# General neural mechanisms can account for rising slope preference in localization of ambiguous sounds

**DOI:** 10.1101/687178

**Authors:** Jean-Hugues Lestang, Dan F. M. Goodman

## Abstract

Sound localization in reverberant environments is a difficult task that human listeners perform effortlessly. Many neural mechanisms have been proposed to account for this behavior. Generally they rely on emphasizing localization information at the onset of the incoming sound while discarding localization cues that arrive later. We modelled several of these mechanisms using neural circuits commonly found in the brain and tested their performance in the context of experiments showing that, in the dominant frequency region for sound localisation, we have a preference for auditory cues arriving during the rising slope of the sound energy (Dietz et al., 2013). We found that both single cell mechanisms (onset and adaptation) and population mechanisms (lateral inhibition) were easily able to reproduce the results across a very wide range of parameter settings. This suggests that sound localization in reverberant environments may not require specialised mechanisms specific to perform that task, but could instead rely on common neural circuits in the brain. This would allow for the possibility of individual differences in learnt strategies or neuronal parameters. This research is fully reproducible, and we made our code available to edit and run online via interactive live notebooks.

## Introduction

The precedence effect, or law of the first wave, is a psychoacoustical phenomenon occurring when multiple sounds reach the listener’s ears in quick succession (Wallach et al., 1949). For small enough delays, the perceived sound is located near the source of the leading sound. In other words, in order to extract reliable localization cues, the auditory system favors the computation of dichotic cues conveyed in the first sound arriving at the ears. Doing so unburdens the localization process from dealing with confounding reverberations. Although well studied (Zurek, 1987; Blauert, 1997; Litovsky et al., 1999; Brown et al., 2015), the neural mechanisms behind the precedence effect are still not well understood. In particular, it is not clear at what stage of the auditory pathway the re-weighting of the interaural cues occurs.

Principally, two main processing sites were suggested. On the one hand, different types of neural adaptation have been reported to take place in the periphery of the auditory pathway. Proposed models of peripheral adaptation range from simulating the ringing of the basilar membrane (Tollin, 1998) to adaptation occurring at the synapse between the inner hair cells and the auditory nerve fibers (Hartung and Trahiotis, 2001; Xia and Shinn-Cunningham, 2011). Similarly, onset cells located in the cochlear nucleus (Rothman and Manis, 2003; Spencer et al., 2012, 2018) are likely to play a role in discarding counfounding localization cues. On the other hand, models of the auditory system including lateral inhibitory circuits in the medial superior olive (MSO) or inferior colliculus (IC) were also able to account for some aspects of the precedence effect. Most of these models rely on the transient inhibition of the lagging sound (Lindemann, 1986a,b; Zurek, 1987; Xia et al., 2010). Overall, peripheral models seem to account well for precedence effect related behaviors with short sounds while inhibition based models produce better results with longer sounds (Braasch and Blauert, 2003).

To further understand how interaural time differences (ITDs) are processed by the auditory system in reverberant environments, a study from Dietz et al. (2013), showed that human listeners tend to rely on interaural phase differences (IPDs) located in the onset of low-frequency amplitude modulated sounds. To demonstrate this, the authors crafted an amplitude modulated binaural beats (AMBB) stimulus exhibiting a time-varying IPD performing a full cycle (from 0° to 360°) over the time-course of a single envelope cycle, while ensuring that there is no interaural level difference. By asking participants to match the perceived location produced by the AMBB sounds to static IPD pointers, Dietz et al. (2013) observed that the subjects mostly made use of the interaural cues positioned in the rising portion of the envelope of the stimulus (instead of the peak of the envelope where the amplitude is maximal). The perceived IPD was also shown to increase with modulation frequency. Later on, Hu et al. (2017) conducted a similar experiment and uncovered that IPD extraction in the rising slope is conditional to the value of the carrier frequency. For 200 Hz stimuli, the subjects mostly reported IPDs located at the peak of the envelope. However, for 600 Hz stimuli, subjects prioritized the rising portion of the envelope.

We investigated a variety of neural mechanisms that could account for the perceived location of the AMBB stimulus in an attempt to rule out certain mechanisms. Instead, we found that all the mechanisms we tried were able to account for this effect. This raises the possibility that the effect may be the result of general processes occurring throughout the brain rather than specialised mechanisms for sound localisation in reverberant environments, and would allow for considerable individual differences in terms of neuronal parameters (Prinz et al., 2004) or localization strategies (Keating et al., 2016). In the first half of this paper, we investigate single neuron mechanisms that occur before binaural integration takes place, using a simple, general model including multiple mechanisms that can enhance onset (including adaptation and delayed inhibition). In the second half of the paper we investigate population level mechanisms based on the interaction of excitation and inhibition after binaural integration. To ensure computational reproducibility, the entirety of the code used to produce the results presented in this paper is available online (https://github.com/neural-reckoning/simple_ambb_modelling) and can be edited and run directly in the browser.

## Results

We investigated the neural mechanisms that could account for the data collected by Mathias Dietz and colleagues on amplitude modulated binaural beats (AMBBs; Dietz et al. 2013, 2014). These are sinusoidally amplitude modulated tones where there is an interaural phase difference (IPD) in the tone or carrier that linearly increases from 0 to 360° during each amplitude modulation cycle. At low modulation frequencies this is perceived as a sound moving around the head. In psychophysical experiments, subjects were asked to move a slider to modify a second amplitude modulated tone with a static IPD (controlled by the slider) to match as closely as possible the AMBB. It might be expected that when forced to assign a single IPD to a stimulus that has all possible IPDs, the IPD at the time where the stimulus was strongest would be chosen (corresponding to an IPD of 180°). However, they found that subjects tended to choose an IPD in the rising portion of the envelope (figure 1), at a phase that increases monotonically with the modulation frequency but is always less than 180°.

**Figure 1:**
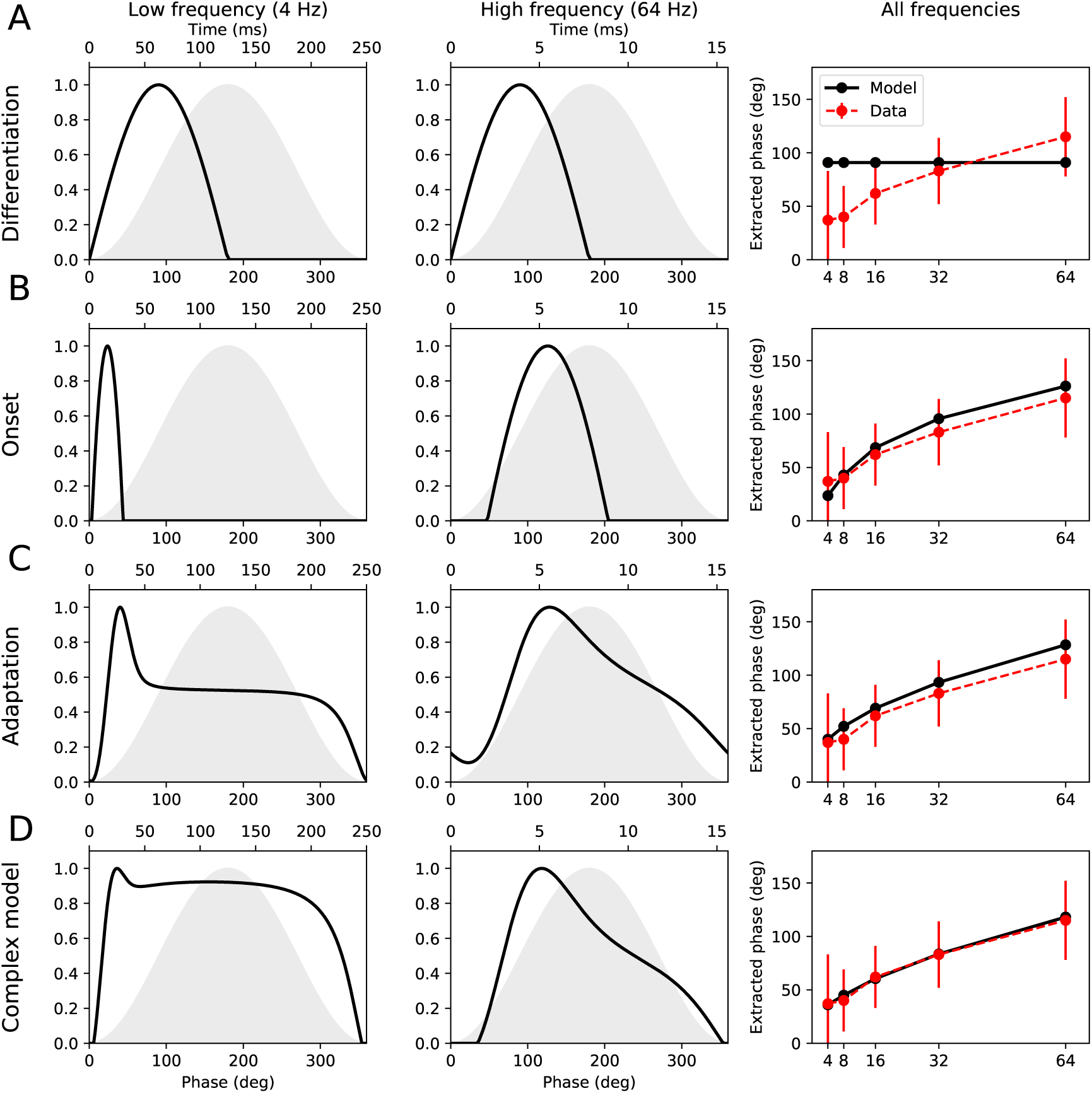
Basic mechanisms of the single neuron model. In each panel, the left and middle plots show, respectively, the response of the model at a low (4 Hz) and high (64 Hz) modulation frequency. The envelope is shown as the grey shaded area, while the output of the model is shown as a black line. The right hand plot shows the location of the peak response of the model at different modulation frequencies (black line), and the data for human subjects (red dashed line). The human data shows the standard deviation of responses across subjects as error bars. (A) Model output is the rectified differential of the envelope. (B) Onset model. Adaptation model. (D) Complex model using both onset and adaptation mechanisms.

We consider two possible explanations for this effect based on mechanisms that occur either before or after binaural integration. In Single neuron mechanisms, we investigate whether monaural single neuron properties could explain the observations, and what conditions the observed data would impose on the parameters and types of those neurons. Dietz et al. (2014) argue that the mechanism must occur before binaural integration takes place, but their argument is based on only a single neuron. With a population of neurons, an inhibitory mechanism can also explain the effect. In Population mechanisms, we investigate these population level effects with a lateral inhibition mechanism. Throughout, we use abstract rate models of neurons with rich dynamics to extract the essential details while keeping the model complexity manageable (similarly to Goodman et al. 2017). This enables us to investigate how the behaviour of the model depends on all of the parameters and plot the parameter spaces, which would not be possible with a biophysically detailed model with a large number of parameters.

### Single neuron mechanisms

A natural place to start given that subjects tend to favour the rising slope would be to assume that the input signal is being differentiated, and therefore the reported IPD will be at the peak of the (differentiated) neural signal (figure 1A). However, this peak occurs at 90° regardless of the modulation frequency, which does not match the pattern of a later preference at higher modulation frequencies observed experimentally, and also leaves open the question of the neural mechanism implementing this differentiation.

#### Onset and adaptation mechanisms can explain rising slope preference

We model onset behaviour abstractly as the weighted difference of two low-pass filtered signals (with different time constants). This can be interpreted either as two precisely timed excitatory/inhibitory pathways, or in a similar way to the octopus cell model of Spencer et al. (2012, 2018) and Ferragamo and Oertel (2002). Their model uses a spike threshold on the rate of change of the membrane potential (Platkiewicz and Brette, 2010, 2011), which we reformulated in terms of firing rates (see Methods and Appendix). With this model it is possible to closely fit the experimentally observed relationship between the modulation frequency and the extracted phase with this model (figure 1B). The parameters are *β ≥* 0 the relative weight of the subtracted signal (so *β* = 0 corresponds to just low pass filtering the signal), and the time constants of the positive and negative filters (*τ*_*e*_ and *τ*_*i*_ respectively).

Another set of mechanisms relate to various forms of adaptation, e.g. spike frequency adaptation or synaptic depression. We model these with a single reservoir model that exhibits a response to a constant input that exponentially decays from an initial to a fully adapted response. This is similar in structure to several well used and equivalent models of adaptation in the auditory nerve (Meddis, 1986; Westerman and Smith, 1988; Zhang and Carney, 2005). It is also equivalent to the short-term synaptic plasticity model of Tsodyks et al. (1998) (with depression only, no facilitation, for the derivation see appendix A of Tsodyks and Wu 2013). Our adaptation model is therefore able to summarise a number of adaptation-related mechanisms, and uses only two parameters (see Methods). Figure 1C shows that this adaptation mechanism is also able to closely fit the data. The parameters are *τ*_*a*_ the time constant of the adaptation, and 0 *≤ α <* 1 the strength of the adaptation (with *α* = 0 indicating no adaptation).

In our full model, both onset and adaptation mechanisms are available, as well as some additional mechanisms such as gain and compression, and this enables us to fit the data even more closely (figure 1D).

#### Mechanisms are highly robust

We computed which values of the model parameters were consistent with the data, and found that the model was very robust, with large regions of the parameter space giving good fits (Figure 2A). Note that since the model has 6 free parameters and there are only 5 data points, it is a priori unsurprising that it is possible to fit the data well. However, this is not a case of simple curve fitting as the model is highly constrained in terms of the possible phase/*f*_*m*_ curves it can produce (Figure 2B), with the majority (73%) being monotonically increasing, for example. We investigate this in more detail in Curve fitting analysis below. Finally, if we remove some mechanisms and their associated parameters, in almost all cases the model is able to fit the data well with as few as three parameters (Figure 3), and a mathematical approximation fits reasonably well with just one parameter (see Appendix).

**Figure 2:**
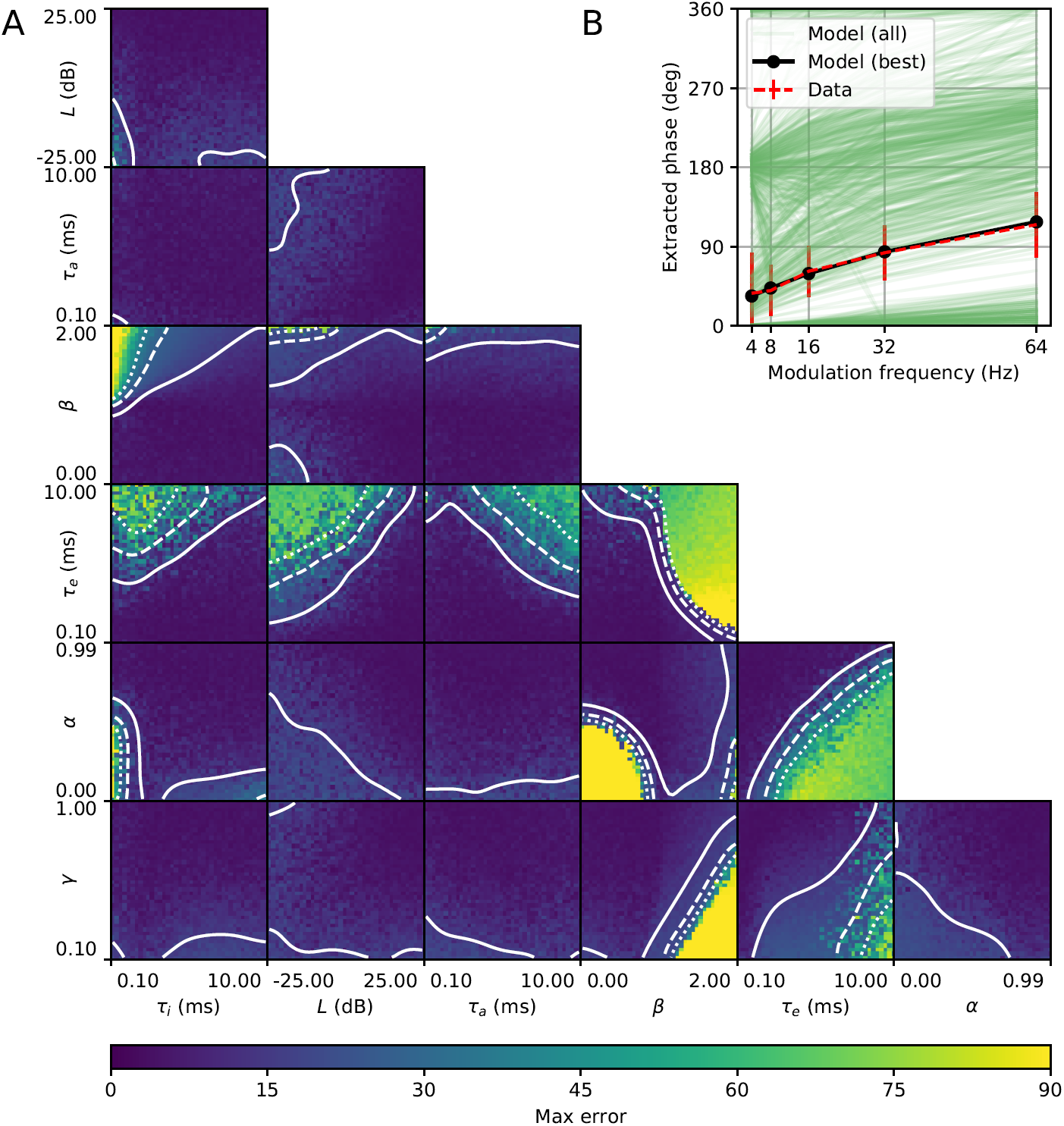
Parameter space. (A) Model error plotted as a function of different pairs of parameters. Each pixel shows the lowest error achievable for a particular fixed value for the corresponding pair of parameters. White contours show 15 degree error (solid), 30 degrees (dashed) and 45 degrees (dotted). (B) A sample of the extracted phase curves for 1000 randomly selected parameters (green curves, excluding parameters which give zero output for some value of *f*_*m*_), the data (red dashed) and the best fitting model (black).

**Figure 3:**
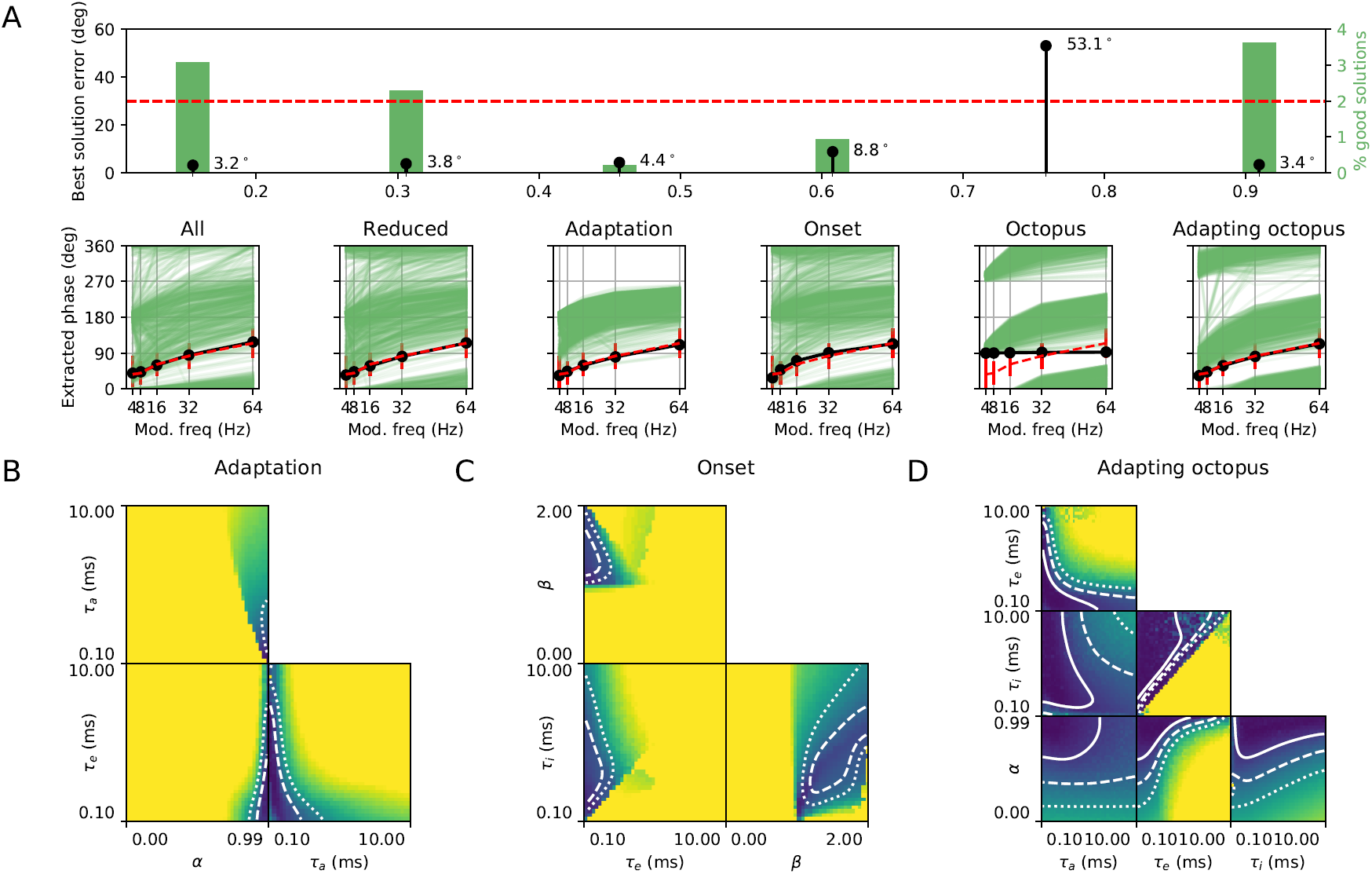
Comparison of restricted models. (A) Extracted phase curves (lower panels as in Figure 2B) for different model variants: *All* parameters available; a *Reduced* set where gain and compression were removed (*L* = 0, *γ* = 1), which is also the case in the remaining variants; *Adaptation* only (*β* = 0); *Onset* only (*α* = 0); *Octopus* cell (*α* = 0, *β* = 1); *Adapting octopus* cell (*β* = 1). The upper panel shows the error of the best performing set of parameters found for that variant. All are within the error bars of the data (dashed red line) except for the octopus cell. (B-D) Parameter maps as in Figure 2A for three of the model variants (adaptation, onset, adapting octopus).

Using this model, we can determine which mechanisms are able to explain the data, and which mechanisms are essential. We considered model variants where some of the parameters were given fixed values (Figure 2A) or some mechanisms removed (Figure 3). The key mechanisms of the model are adaptation and onset: either mechanism can work, although the best fits obtained with adaptation only are better than the best fits obtained with the onset mechanism only (Figure 3ABC); and at least one of the two mechanisms must be present (Figure 2A, *α* versus *β*). The compression and gain mechanisms did not contribute substantially (Figure 3A). We investigated the octopus cell model discussed earlier (Ferragamo and Oertel, 2002; Spencer et al., 2012, 2018) and found that it could provide a good fit only if adaptation was present alongside the onset mechanism (Figure 3AD). For model variants including the onset mechanism, the positive or excitatory time constant *τ*_*e*_ must be small (Figures 2A and 4B), and the negative or inhibitory time constant *τ*_*i*_ must be larger than *τ*_*e*_ (Figure 3CD, *τ*_*e*_ versus *τ*_*i*_). An interesting feature of all the model variants is that the extracted phase curve in the experimental data appears to be at or close to the lowest values possible in the models (Figure 3A).

#### Best model fits divide into two main phenomenological cell types

We investigated the parameters and phenomenological behaviour of all model cells that were capable of reproducing the experimental data within an error of 30° (Figure 4), as this was the size of the smallest error bar in the experimental data. We computed the rate and temporal modulation transfer functions (rMTF, tMTF), and extracted the corresponding best modulation frequencies (BMF), mean MTFs, and modulation depths (MD) for each cell. We found that there was no clear demarcation of the parameter space into discrete cell types based on parameter values, as a continuum of values led to good fits (Figure 4A). However, we found that there was a clear demarcation into two cell types based on their modulation transfer functions. One cell type has a high tMTF: it phase locks sharply to the stimulus envelope at all modulation frequencies (high mean tMTF value, low tMD), and has a strongly modulated high pass or band-pass rMTF. This cell type corresponds to the parameter *β >* 1, or a strong onset mechanism which allows it to strongly phase lock to the envelope. These cells can have a wide range of adaptation strengths, fast excitatory time constants and slower inhibitory time constants. The other cell type has a more variable tMTF with a correspondingly higher tMD, along with a typically lower rMD. This cell type corresponds to the parameter *β <* 1, or a weak onset mechanism that is consistent with a weaker locking to the envelope. These cells tend to have stronger adaptation, and are consistent with a wider range of excitatory and inhibitory time constants compared to the onset cell type, as well as a wider range of behaviours (Figure 4B). Both cell types were about equally present in the parameter space considered.

**Figure 4:**
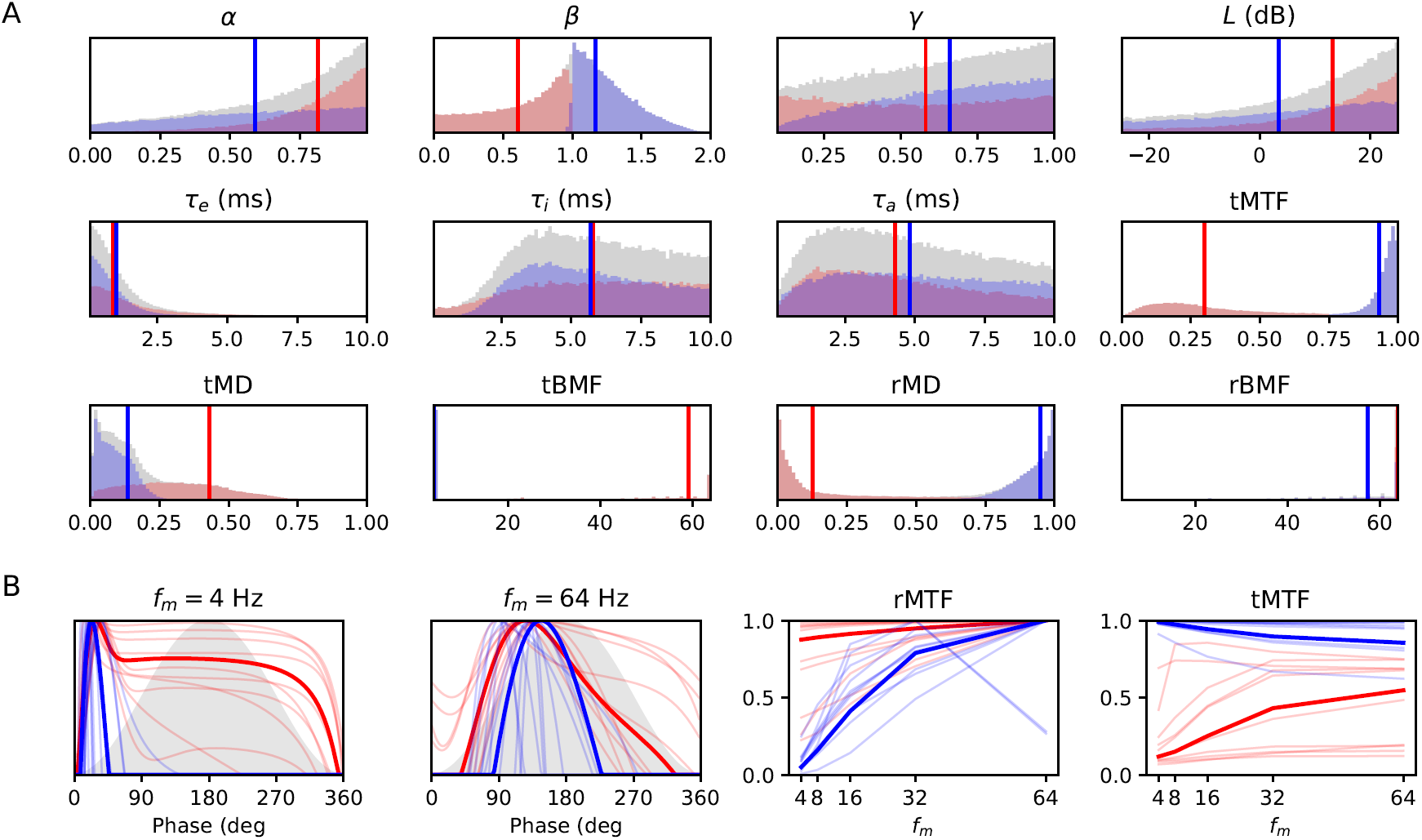
Cell types. (A) Histograms for all parameters and properties leading to an error of less than 30° (grey), for the strong onset cell class with tMTF*>* 0.75 (blue), or the weak onset cell class with tMTF*<* 0.75 (red). Vertical lines show mean values for the two classes. (B) Sample responses from the two cell classes. Left two panels as in Figure 1, right two panels show detailed rate and temporal modulation transfer functions. Thin lines show a random sample of curves from each class, and thick lines show the most representative example from that class (smallest distance to all other points in the cell class parameter space).

#### Homogeneous neurons without onset mechanisms do not exhibit an early preference

So far, we have considered a model that only responds to the envelope of the sound, which can be seen either as an approximation, or as applying only to cells with low synchronisation to the carrier. We added the carrier to the model, and one additional low pass filter mechanism (representing the inner hair cell or other low pass filtering processes). For simplicity we used a low pass filter with time constant *τ*^IHC^, although higher order filters may provide a better fit to the data (Russell and Sellick, 1983). In addition to considering the carrier frequency of *f*_*c*_ = 500 Hz used in Dietz et al. (2013, 2014), we also calculated results for *f*_*c*_ = 200 Hz. This lower carrier frequency was studied in Hu et al. (2017) with a similar experimental design, and they found that there didn’t appear to be an extraction of the early IPD at this *f*_*c*_, only at the higher *f*_*c*_ (they used 600 Hz in that paper, similar to the 500 Hz used in the earlier papers). However, the results are not directly comparable, so we decided to measure the error in the model as the combination of the error at 500 Hz as measured above, and an error at 200 Hz assuming that the equivalent experimental results were a flat IPD = 180°. The contribution of the error at 200 Hz was weighted less as there is no direct experimental data in this case, only a hypothetical curve based on a similar experiment (see Methods for more details).

We found that the model was able to simultaneously reproduce the two different extracted phase curves at *f*_*c*_ = 200, 500 Hz (Figure 5). The main difference in the parameters found is that with the carrier frequency included in the model it is not possible to closely fit the data with an adaptation-only model (*β* = 0), and indeed onset strength has to be quite high (*β >* 1, Figure 5B). In addition, some well fitting parameters in the envelope only model with a large *τ*_*i*_ no longer work well in this case. We found that uniformly the tMTF was higher at *f*_*c*_ = 500 Hz compared to 200 Hz, the tMD lower, and the rMD was always very high for *f*_*c*_ = 500 Hz (but could have a wide range of values at 200 Hz).

**Figure 5:**
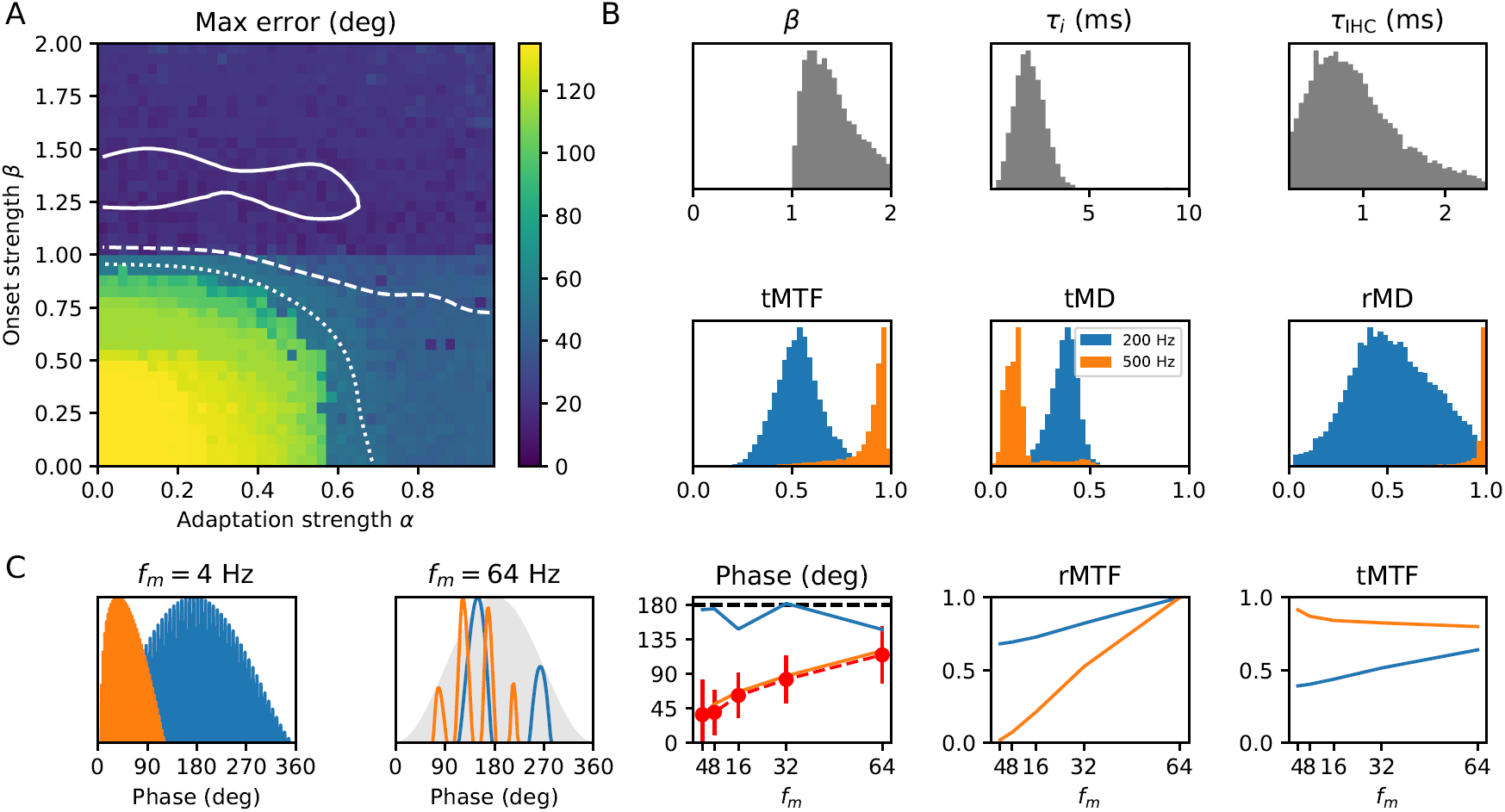
Effect of introducing carrier frequency. (A) Error in best fit for fixed adaptation strength *α* and inhibition strength *β* (as in Figure 4). Here, the error is measured at both *f*_*c*_ = 500 Hz and *f*_*c*_ = 200 Hz assuming a hypothetical flat 180° extracted in the latter case. The 200 Hz error is weighted less as there is no corresponding experimental data. (B) Parameter and property histograms as in Figure 4. Here, colours show properties at 200 Hz (blue) and 500 Hz (orange). (C) Best fit in detail as in figure 4.

It would be tempting to conclude from this that only onset cells can account for all the experimental data of Dietz et al. (2013, 2014) and Hu et al. (2017), however the populations of cells responding to a 200 Hz and 500 Hz carrier frequency are different, and there is no a priori reason to think that they should be the same cell types or have the same parameters.

#### Model predicts earlier preference at higher sound levels

We took the collection of good parameters (with an error less than 30°) and applied different gains to amplify or attenuate the signal before recomputing the extracted phase curves. We found that with a considerable degree of consistency, increasing the sound level led to an earlier phase being extracted, while decreasing the level led to a later phase (Figure 6). Experimental data has not yet been collected to quantitatively test this hypothesis, but early observations are in qualitative agreement (Mathias Dietz, personal communication of unpublished observations).

**Figure 6:**
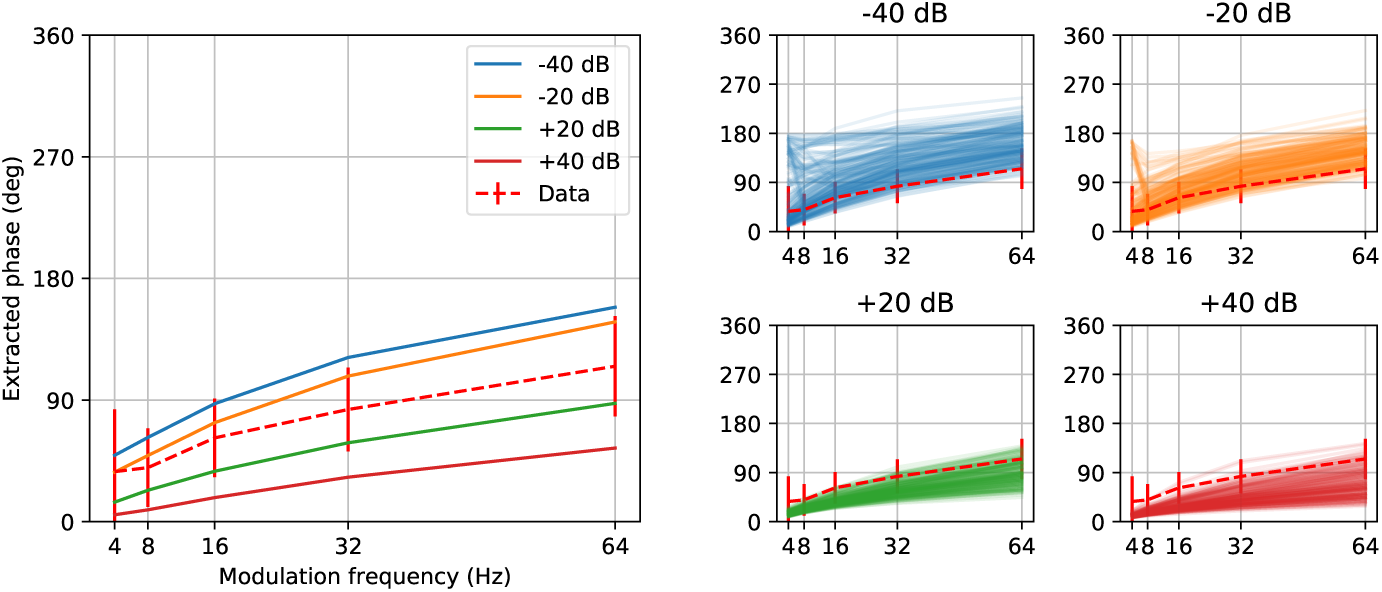
Single neuron model predictions for varying sound level. Left panel shows mean prediction at different sound levels for parameters that match the experimental data well at a sound level of +0 dB. Right panels show sample extracted phase curves for each sound level.

### Population mechanisms

In order to investigate whether mechanisms acting after binaural integration could give rise to a similar effect, we next investigated the role of population mechanisms assuming no monaural mechanisms that give rise to an early phase preference. In the experiments of Dietz et al. (2013) participants were asked to choose which amplitude modulated tone with a static IPD sounded most similar to the amplitude modulated binaural beat (AMBB) stimulus. We therefore modelled the process as pattern matching on the neural population (as in the sound localisation model of Goodman et al. 2013). We included a lateral inhibition mechanism that is found throughout the brain, and in the case of sounds with a static IPD serves to better separate the patterns. In the case of the AMBB stimulus, this model was able to robustly reproduce the preference for early phase, getting later with increasing modulation frequency, and fit the experimental data with low error across a wide range of parameters (figure 7).

**Figure 7:**
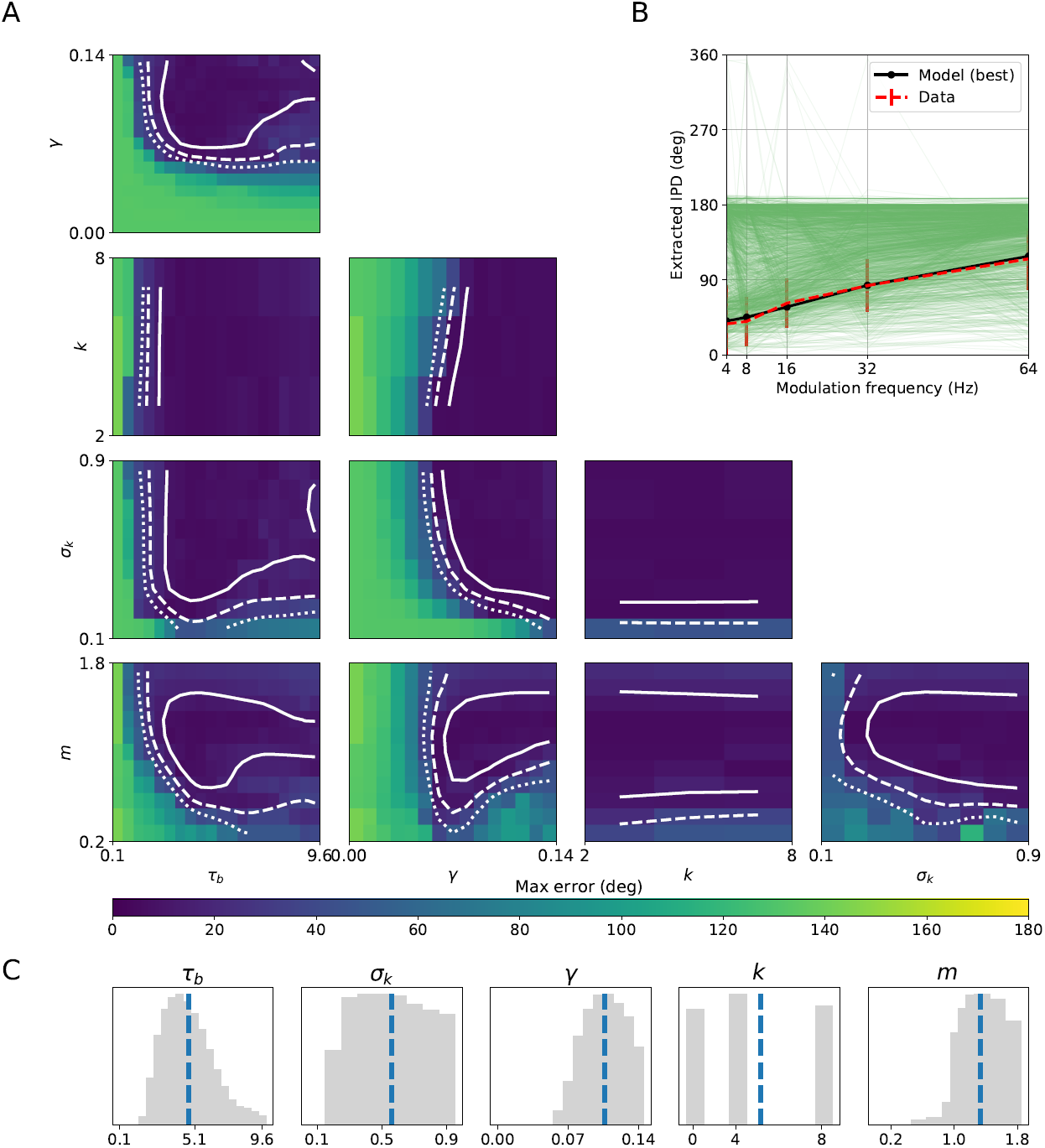
Population model parameter space, as in figure 2. (A) Model error plotted as a function of different pairs of parameters. Each pixel shows the lowest error achievable for a particular fixed value for the corresponding pair of parameters. White contours show 15° error (solid), 30° (dashed) and 45° (dotted). (B) Sample and best model results. (C) Histograms of the parameters leading to an error less than 30°.

In more detail, the model receives as input responses from a population of cells with different IPD tuning, with the density of best IPDs as measured for guinea pigs (McAlpine et al., 2001). These neurons directly excite a second layer of neurons and inhibit the neighbours of those neurons, serving to sharpen the pattern of responses. Finally, we compare the patterns of activities for the AMBB stimulus with the patterns obtained for different amplitude modulated tones with static IPDs, and select the static IPD with the most similar response. The model has five parameters: *τ*_*b*_ is the time constant of the inhibition process; *σ*_*k*_ is the width of the lateral inhibition as a fraction of the whole population (so *σ*_*k*_ = 1 means all neurons inhibit all others and *σ*_*k*_ = 0 means no inhibition); *γ* is the strength of the inhibition; *k* is an integer that determines the shape of the IPD tuning curve and can be experimentally fit to different animal models; and *m* is the envelope synchronization of the inputs, so when *m* = 0 the inputs are not synchronised to the envelope at all, at *m* = 1 they exactly match the shape of the envelope, and for *m >* 1 they have enhanced synchronisation (e.g. onset cells). See Methods for further details of the model.

#### Lateral inhibition and pattern matching gives rise to a robust early phase preference

We calculated model fits for a wide range of parameter values (89,100 different parameter sets in total). The best fit had an error of just 5° (figure 7B), and large regions of the parameter space led to low errors (figure 7A) of less than 30° (the size of the smallest error bar in the experimental data). Overall, 8.1% of the parameter sets led to an error below 30° and 41% of the parameter sets led to solutions which increased monotonically with the modulation frequency. Most of the parameter sets (78%) led to a later phase preference than the experimental data, 2% to an earlier phase preference and 20% were earlier for some modulation frequencies and later for others.

Some parameters are more tightly constrained than others if we seek a close fit to the data. The synchronization index *m* for parameter sets with low error shows a particularly striking pattern (figure 7C). There is a very steep drop in the number of good solutions where *m <* 1, suggesting that enhanced synchronisation may play a role. This enhanced synchronisation is evident in some auditory neurons, and may come about as a result of the onset mechanisms discussed in Single neuron mechanisms for example. For parameters *τ*_*b*_ and *γ* the best fits were obtained in a limited although not extreme range. Best fits were obtained for *τ*_*b*_ in the moderate range 2-8 ms, for example. Larger values of the inhibition strength *γ* lead to a shut down of the network and less inhibition leads to solutions closer to 180°. The least critical parameters were *σ*_*k*_ and *k*. The former is the width of the lateral inhibition, which could take on almost any value except those close to the extremes where there was no inhibition (*σ*_*k*_ = 0) or maximum inhibition (*σ*_*k*_ = 1). The tuning curve width, controlled by *k*, had almost no effect.

#### Distribution of tuning leads to peak preference at low carrier frequencies

Hu et al. (2017) suggests that carrier frequency has an important effect on the perceived IPD of the AMBB stimulus. They showed that with a carrier frequency of 200 Hz, the perceived IPD is shifted toward the peak of the envelope, while at 600 Hz it is perceived during the rising slope as in Dietz et al. (2013). McAlpine et al. (2001) found that the distribution of best IPDs at 200 Hz is different to that at 500 Hz. In our model, we found that for some parameters, this change in BIPD distribution alone caused a shift in perceived IPD towards the peak (figure 8). As before, when investigating model fits at 200 Hz and 500 Hz we weight the error at 200 Hz much less (one third) than the error at 500 Hz, as the experimental data from Hu et al. (2017) is much less precise (preferred IPD is only reported as rising, peak or falling, a granularity of about 90°). The lowest error found was approximately 15°, meaning that the extracted phase was within 15° for all modulation frequencies at 500 Hz, and within 45° at 200 Hz, both well within the limits set by the data. In order to interpret the sensitivity of the model to the BIPD distribution, we computed the mean preferred IPDs across all good solutions (error below 30°) for carrier frequencies from 200 to 1000 Hz. We found that decreasing the carrier frequency led to a shift in preference towards the peak (figure 8D). For carrier frequencies above 500 Hz, the preference did not shift any earlier.

**Figure 8:**
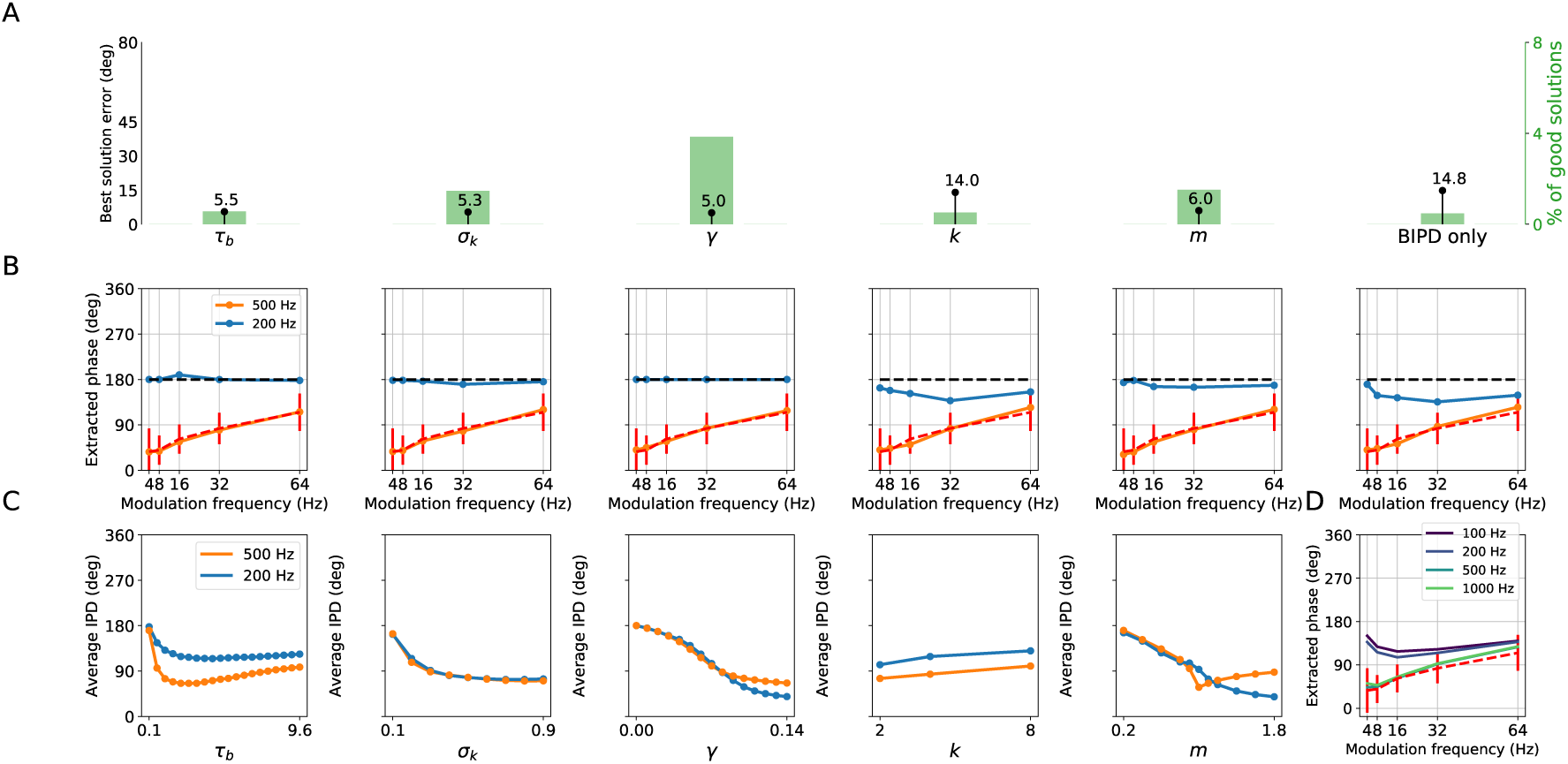
(A) Error in best solution (black lines and dots), and percent of the parameter space leading to good solutions (green bars), when allowing one parameter to take different values at 200 Hz and 500 Hz (except for “BIPD only” where all parameters are identical except for the BIPD distribution). (B) Best solutions found at 200 Hz and 500 Hz. (C) Trend in mean preferred IPD over modulation frequencies when varying the value of a single parameter. The trend was obtained by averaging over all solutions below 30°. Mean IPD across all good solutions for various carrier frequencies.

Although the error for the best solution is low, this result requires some precise parameter tuning, and only 0.5% of parameters tested had an error lower than 30°. This model assumes that the parameters of the neurons must be the same at a carrier frequency of 200 and 500 Hz, but these are different neurons (due to tonotopy, Humphries et al. 2010; Ress and Chandrasekaran 2013), and they may therefore have different parameters. We refer to the case where all parameters must be the same as homogeneous and the case where they may be different as heterogeneous. We investigated heterogeneous networks by allowing only one parameter to have a different value at 200 and 500 Hz (figure 8). Unsurprisingly, this leads to much better fits. Allowing for different relative strengths of excitation and inhibition *γ* at 200 and 500 Hz – which is very plausible as synaptic weights can be learned – leads to a best fit with an error of just 5°, and almost 4% of the parameter space with errors less than 30°. Allowing *σ*_*k*_ and *m* to vary led to only slight larger minimum errors, although in these cases a somewhat smaller percentage of parameters led to errors below 30°. Allowing the tuning curve width to vary (controlled by *k*) did not contribute to a good fit. We investigated how each parameter contributes to the mean preferred IPD (calculated across all good solutions and all modulation frequencies, figure 8C). Preferred phase decreased monotonically when either the width (*σ*_*k*_) or strength of inhibition (*γ*) increased, and tended to decrease with increasing synchronisation (*m*), except at 500 Hz for larger values of *m*. Very small time constants of inhibition (*τ*_*b*_) led to a later phase preference, but had relatively little effect as long as the time constant was larger than around 1 ms. Narrower tuning curves (larger values of *k*) led to slightly later phase preference.

### Curve fitting analysis

For both the single neuron and population models, we have around the same number of parameters as data points. We therefore investigated whether or not our results could be explained simply as curve fitting by calculating how well the model could fit the data if the data were different (figure 9). Two different conditions were used. In the first one, the target IPDs for each modulation frequency were chosen independently and uniformly at random between 0° and 360°. In the second condition, the target IPDs were again chosen randomly, but this time between 0° and 180°, and with the additional constraint that they should be monotonically increasing with modulation frequency. In both cases, the best fits obtained to the actual data were much better than the best fits for the random target IPDs in almost all cases. None of the 1,000 fully random IPD targets could be fitted as well as the experimental data. With the extra constraints, the single neuron model could fit the random data better than the experimental data only 3% of the time, and the population model only 8% of the time. Since the models were not able to fit the majority of random data, even when constrained to have a qualitatively similar shape to the experimental data, these results cannot simply be explained as curve fitting.

**Figure 9:**
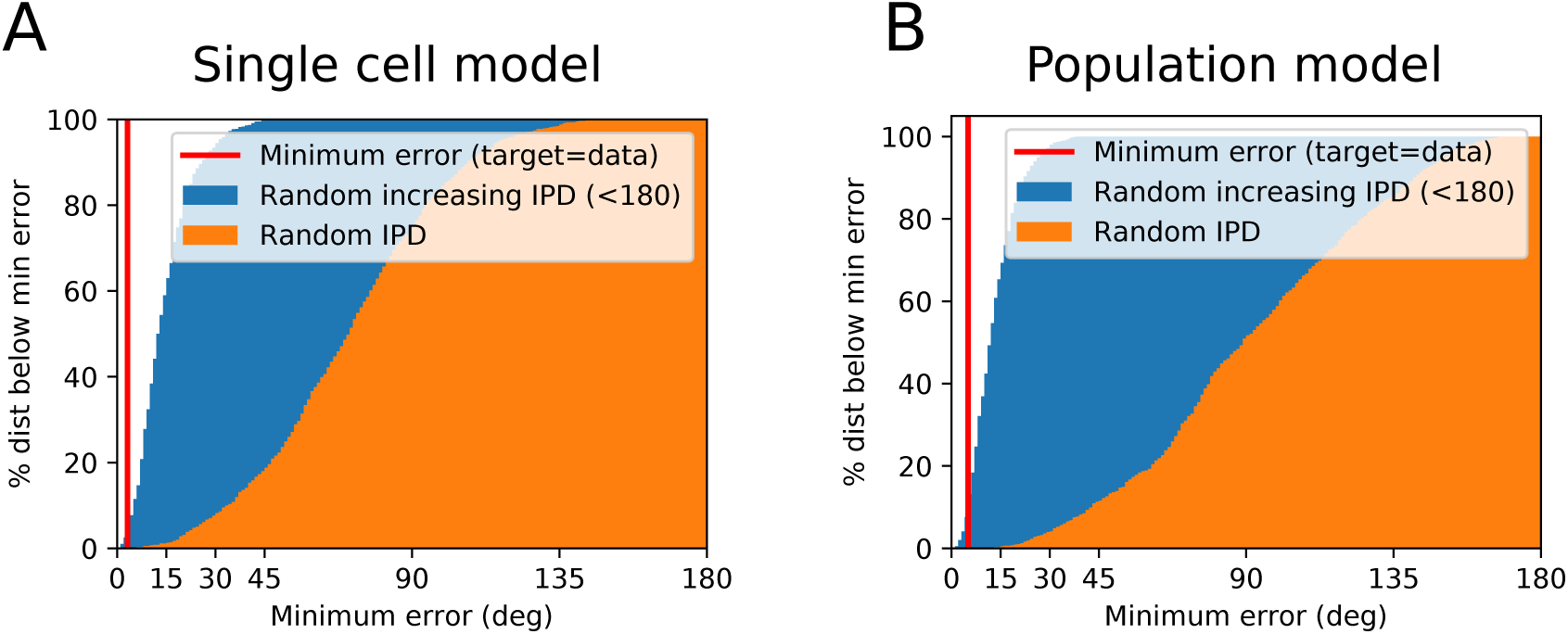
Best fits for experimental data (red line) and randomised target data (orange for fully random, blue when constrained to be monotonically increasing with modulation frequency less than 180°). Shaded areas show cumulative histograms of the minimum error found for the target data (so 0% of randomly selected data could be fit with an error of 0° while 100% could be fit with an error less than 180°). (A) Single cell model. (B) Population model.

## Discussion

We modelled a number of single neuron and population level neural mechanisms across different stages of the auditory pathway to account for the preference for extracting binaural cues during the rising slope of an amplitude modulated signal (Dietz et al., 2013). By emphasizing the cues contained in the onsets of sounds, this effect was suggested to be important in responding to the sound following a direct path in a reverberant environment, and discarding later arriving information from indirect paths. This may be an important factor underlying a number of phenomena, including spatial release from masking in cocktail party situations (Cherry, 1953; Freyman et al., 1999).

We found that adaptation and onset mechanisms occurring in single neurons before binaural integration (for example in the auditory nerve or cochlear nucleus) could lead to a rising slope preference that was an excellent quantitative fit to the experimental data recorded by Dietz et al. (2013) across a very wide range of parameters. The best results fell into two main separate classes: strong onset or strong adaptation. Of these two, onset cells were more consistent with the result that preference shifts towards the peak when the carrier frequency is low (Hu et al., 2017), although this did not allow us to fully rule out adaptation as a key mechanism as the auditory system is tonotopically organised and neuron parameters in low frequency regions may be different to those at higher frequencies. Our model predicts that higher sound levels lead to a preference for earlier onsets (and vice versa), which is consistent with preliminary unpublished observations (Dietz, personal communication).

We then considered mechanisms arising at the level of populations of neurons after binaural integration (for example in the inferior colliculus or auditory cortex). We found that we could again very precisely reproduce the quantitative effect, this time assuming only a local interaction of excitation and inhibition, a lateral inhibition motif that is found in many areas of the brain. This model was also very robust, and was able to reproduce the shift towards the peak at low frequencies based only on the difference in distribution of the interaural phase difference preferences of binaural neurons observed in mammals (McAlpine et al., 2001). This model required that its inputs show a strong synchronisation to the envelope of the stimulus, which can be the result of the onset mechanisms discussed above for example, suggesting that single neuron and population mechanisms may work in concert. The model finally predicts that although lowering the carrier frequency below 500 Hz leads to a later preference (a shift towards the peak), increasing the carrier frequency above 500 Hz should not lead to an earlier preference.

Although some of the mechanisms reviewed in this study have been shown to explain some aspects of the precedence effect (Hartung and Trahiotis, 2001; Braasch and Blauert, 2003; Xia and Shinn-Cunningham, 2011; Xia et al., 2010), none of them were previously tested quantitatively in the context of the experiments by Dietz et al. (2013). Additionally, the use of simple models allows for a comprehensive exploration of some aspects of the precedence effect such as the extraction of IPDs in low-frequency amplitude modulated sounds.

The mechanisms we used in these models are not unique to the auditory system. Our adaptation and onset models are similar or equivalent to membrane potential and synapse dynamics that are found throughout the nervous system (see Methods and Appendix). Adaptation allows neurons to minimize energy costs while encoding persistent stimuli (Jones et al., 2015) and to increase their sensitivity to new stimuli (Fairhall et al., 2001). At the population level, excitatory-inhibitory networks are often found in other sensory modalities, such as in orientation tuning in vision (Ben-Yishai et al., 1995) or in the detection of touch in the field of somatosensation (Mountcastle, 1959). In the case of the auditory system, these circuits have not been positively identified, but would likely be located in the inferior colliculus, where many inhibition circuits exist (Pollak et al., 2011), or higher up in the auditory pathway.

In order to thoroughly investigate the parameter spaces of our models, we abstracted and simplified them as much as possible. This has the great advantage of allowing us to precisely understand the interactions of the different mechanisms, but has the disadvantage of making it difficult to apply the model to other more complex stimuli. Incorporating additional mechanisms such as cochlear and modulation filterbanks would be straightforward but would introduce a large number of additional parameters.

In summary, we found that we could account for the experimental data across a wide range of parameters using multiple single neuron or neural population mechanisms that are not specific to the binaural system, or even to the auditory system. Indeed, it appears that a diverse set of neural mechanisms located in the auditory pathway all emphasize the extraction of the ITD information in the onset of the envelope. While we cannot definitively say which mechanisms are present and contribute to the precedence effect, our results suggest that all the mechanisms we studied have the capacity to do so. This opens the door for two hypotheses that may drive future work.

The first hypothesis is that the preference for early arriving auditory cues (Dietz et al., 2013, 2014; Hu et al., 2017), and the precedence effect more generally, may not be the result of an adaptation specifically for sound localisation in reverberant environments, but may be the result of an evolutionary exaptation or co-opted adaptation. That is, an instance of the brain using already existing dynamics and mechanisms (adaptation, excitation/inhibition, lateral inhibition) to carry out a new function (that may then subsequently have been refined by further evolutionary pressure). This possibility has been suggested more generally under various names, including circuit motifs (Braganza and Beck, 2018), computational primitives (Marcus et al., 2014) and canonical computations (Kouh and Poggio, 2008). In terms of our efforts to try to understand the incredible and general abilities of the brain, this might be an even more exciting possibility than the alternative that there are mechanisms adapted specifically for processing reverberant sounds.

The second hypothesis is that, since the range of parameters equally able to account for the data is so wide, there may be individual differences. These individual differences could come from two sources. It could be that there are a range of neural parameters that all give rise to the same behaviour, as in the case of the pyloric network of the crustacean stomatogastric ganglion (Prinz et al., 2004). Or, it could be that individuals learn different sound localization strategies, making use of the different neural mechanisms that are available to them, in line with the finding that human listeners exhibiting asymmetric hearing can take advantage of multiple adaptive mechanisms to improve their ability to localize sounds (Keating et al., 2016).

We therefore strongly encourage researchers to investigate the extent to which new experimental results can be reproduced using simple and general models, and thoroughly explore the parameter space in detail along the lines suggested by O’Leary et al. (2015) and used previously in the auditory system in Goodman et al. (2017).

## Methods

Models were implemented in Python using the Brian simulator (Goodman and Brette, 2008, 2013; Stimberg et al., 2019) and the scientific computing packages NumPy and SciPy (Jones et al., 2016). All the code is available online (https://github.com/neural-reckoning/simple_ambb_modelling) and can be edited and run directly in the browser via live, interactive notebooks.

### Stimulus

The amplitude modulated binaural beats (AMBB) stimuli, designed by Dietz et al. (2013), are amplitude modulated tones whose envelopes follow the equation

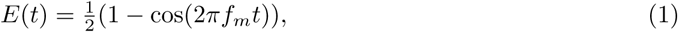

where *f*_*m*_ is the modulation frequency of the stimulus. The main characteristic of the AMBB stimulus is its dynamic interaural phase difference (IPD) varying from 0° to 360° in one modulation cycle. This is achieved by introducing an interaural frequency difference between the left and right channels of the sound:

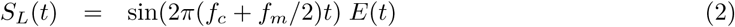

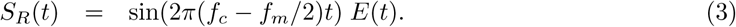

In these equations, *f*_*c*_ refers to the carrier frequency while *S*_*L*_ and *S*_*R*_ represent respectively the left and right channels. In the first part of section Results (up until section Homogeneous neurons without onset mechanisms do not exhibit an early preference) we study peripheral adaptation mechanisms which are known to be monaural, and therefore only use the envelope of the stimulus as input. Later, while investigating the influence of the carrier frequency (section Homogeneous neurons without onset mechanisms do not exhibit an early preference) we add a carrier channel at 500 Hz.

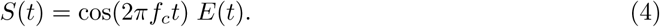

### Single neuron model

We start off by describing the modelling of single cells in the auditory periphery. In this study, cochlear filtering was not necessary as all signals were narrowband. We model amplification, compression and inner hair cell dynamics in a fairly standard way as follows. First we half-wave rectify and compress the signal as

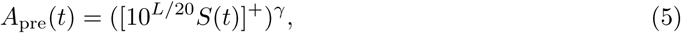

where the operator []^+^ represents half-wave rectification ([*x*]^+^ = *x* when *x >* 0 otherwise [*x*]^+^ = 0), *L* is the gain (in dB) and *γ* is the compression (*γ* = 1 is no compression, and *γ* = 1/3 is a commonly used value). The signal is then passed through a first order low pass filter with time constant *τ*_ihc_:

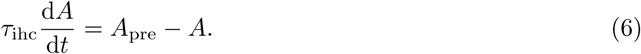

The resulting signal was then passed to an adaptation stage inspired by the functioning of the synapse between the inner hair cells and the auditory nerve fibers. This model is a simplification of published adaptation models (Meddis, 1986, 1988; Westerman and Smith, 1988; Zhang and Carney, 2005) to a single reservoir and consequently a single time-constant. The quantity of available neurotransmitter *Q* (between 0 and 1) is depleted at a rate proportional to the product of the incoming signal and the remaining amount of neurotransmitter (*κQA*), and replenished at a rate proportional to the deficit (*ρ*(1*−Q*)). The output of the model *R*_*a*_ is the product of the input signal A with the instantaneous quantity of available neurotransmitters Q and models the resulting firing rate of the nerve fibers:

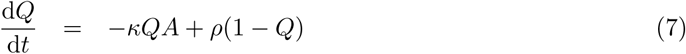

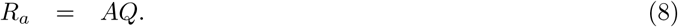

To give an idea of the behaviour of this model, with a fixed input *A* = 1 it decays exponentially to a value 1 *− α* = *ρ/*(*κ* + *ρ*) with time constant *τ*_*a*_ = 1/(*κ* + *ρ*). We can invert this (*κ* = *α/τ*_*a*_, *ρ* = (1 *− α*)*/τ*_*a*_) and use *α* and *τ*_*a*_ as parameters, with *α* representing the strength of adaptation (*α* = 0 no adaptation, *α* close to 1 for strong adaptation) and *τ*_*a*_ the time constant of adaptation (for a fixed reference signal).

Finally, we use the following equations to model either onset cells or excitatory and inhibitory dynamics:

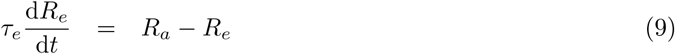

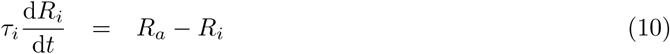

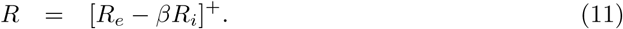

Each current was produced by low-pass filtering the signal with different time-constants *τ*_*e*_ and *τ*_*i*_. If *τ*_*e*_ *< τ*^*i*^ are close this acts as a differentiator.

The model is run for a number of cycles (at least 1) until it settles into a periodic state, and then the IPD returned by the model is the phase corresponding to the time when the output signal *R*(*t*) is at a maximum on the next cycle:

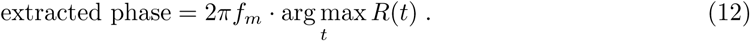

Without any additional mechanisms (adaptation or onset, i.e. *α* = *β* = 0), the IPD returned by the model is consistently equal to 180°.

#### Error measures

To evaluate the fit of the model to the data, we used the maximum absolute error in the IPD across modulation frequencies:

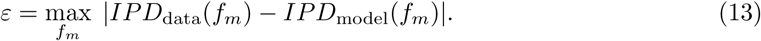

The same error measure was also used in the population model. In cases where we compare errors at *f*_*c*_ = 200 Hz and *f*_*c*_ = 500 Hz we compute *ε*_200_ and *ε*_500_ using the equation above, and compute a total error based on a weighted maximum of these two:

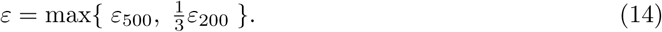

We weight the error at *f*_*c*_ = 200 Hz lower as we do not have direct and equivalent experimental data at this carrier frequency (Hu et al., 2017).

#### Modulation measures

We computed several measures of the variation of phase-locking and firing rates with modulation frequency.

The mean *temporal modulation transfer function (tMTF)* measures the degree to which the model phase-locks to the envelope across modulation frequencies, and is computed as:

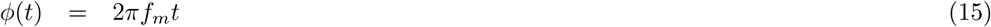

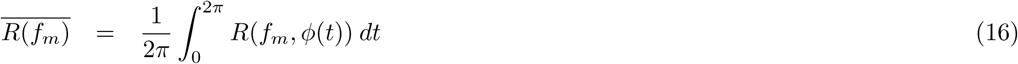

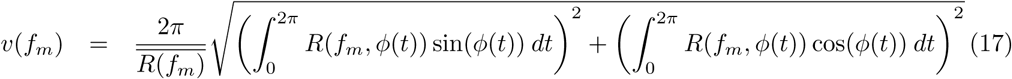

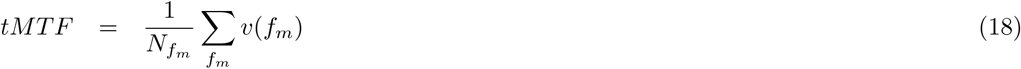

Here *ϕ*(*t*) is the modulation phase at time *t*, 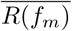 is the mean firing rate across one modulation cycle, and *v*(*f*_*m*_) is the vector strength.

The *rate modulation transfer function, rMTF* (*f*_*m*_), is the ratio of the mean firing rate at frequency *f*_*m*_ to the maximum across all *f*_*m*_, and the mean rMTF is the mean of this over all *f*_*m*_, so that a low value indicates a variable rMTF and a value near 1 indicates a fixed rMTF:

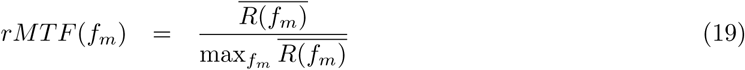

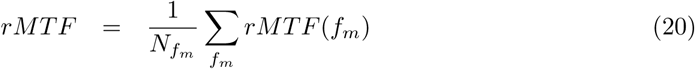

In the set of equations above and in the rest of this method section, a line over a symbol represents the operation of taking a mean over time *t*.

The *temporal modulation depth (tMD)* is computed as the normalised difference between the minimum and maximum vector strength across modulation frequencies:

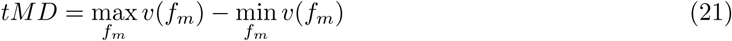

The *rate modulation depth (RMD)* is similarly computed as the normalised difference between the minimum and maximum firing rates across modulation frequencies:

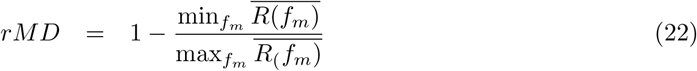

The *temporal best modulation frequency (tBMF)* is the modulation frequency maximizing the vector strength:

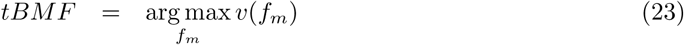

The *rate best modulation frequency (rBMF)* is the modulation frequency maximizing the mean firing rate:

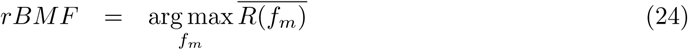

### Population model

We model the experiment of Dietz et al. (2013) by comparing the similarity of the neural population response to the AMBB stimulus to the response to an amplitude modulated tone with a static IPD. The neurons in these populations are binaural and IPD-tuned, and interact with each other by lateral inhibition. We explain each of these elements below.

Each of the binaural neurons in the first layer is sensitive to IPD (*ϕ*), with a tuning curve centered around its best IPD (BIPD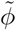) given by

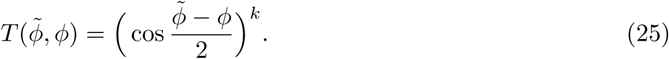

This tuning curve has previously been shown to fit various animal models for different values of *k* (Harper and McAlpine, 2004; Joris et al., 2006; Fischer et al., 2008; Goodman et al., 2013) and is shown in figure 10C. We next assume that the response is proportional to the stimulus amplitude. For the AMBB stimulus, the amplitude at time *t* is given by *E*(*t*) as above, and the IPD *ϕ*(*t*) = 2*πf*_*m*_*t*, so the response of the neuron with BIPD 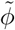 at time *t* is

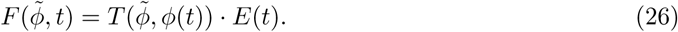

The original amplitude modulated signal has modulation depth 1 (the envelope goes from 0 to 1), however the modulation depth of the neural response to this stimulus may differ, and may have a higher or lower synchronisation or vector strength with respect to the modulation frequency. We model this by replacing the envelope *E*(*t*) above with

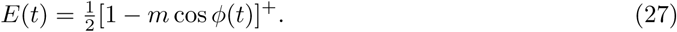

The variable *m* represents the synchronization of the neural response to the envelope of the stimulus. When *m* = 0 there is no synchronization of the neural response to the stimulus. When *m* = 1 the synchronization is the same as the modulation depth of the stimulus. Finally, when *m >* 1, the neuron exhibits enhanced synchronization (as seen for example in onset cells).

For computational efficiency, we discretize the model by choosing *N*^BIPD^ = 100 BIPDs uniformly between 0° and 360°, and (for each modulation frequency) *N*^*t*^ = 250 time steps per modulation cycle (so the sample width is 1*/N*_*t*_*f*_*m*_ at modulation frequency *f*_*m*_). We write the discrete BIPDs and times as 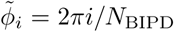 and *t*_*j*_ = *j/N*_*t*_*f*_*m*_. Note that this does not mean we consider BIPDs to be necessarily uniformly distributed: below we will weight the responses according to the density distribution of BIPDs. With these discretizations, we can now write the model in matrix form as follows:

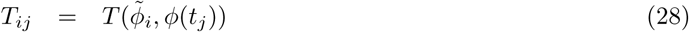

**Figure 10:**
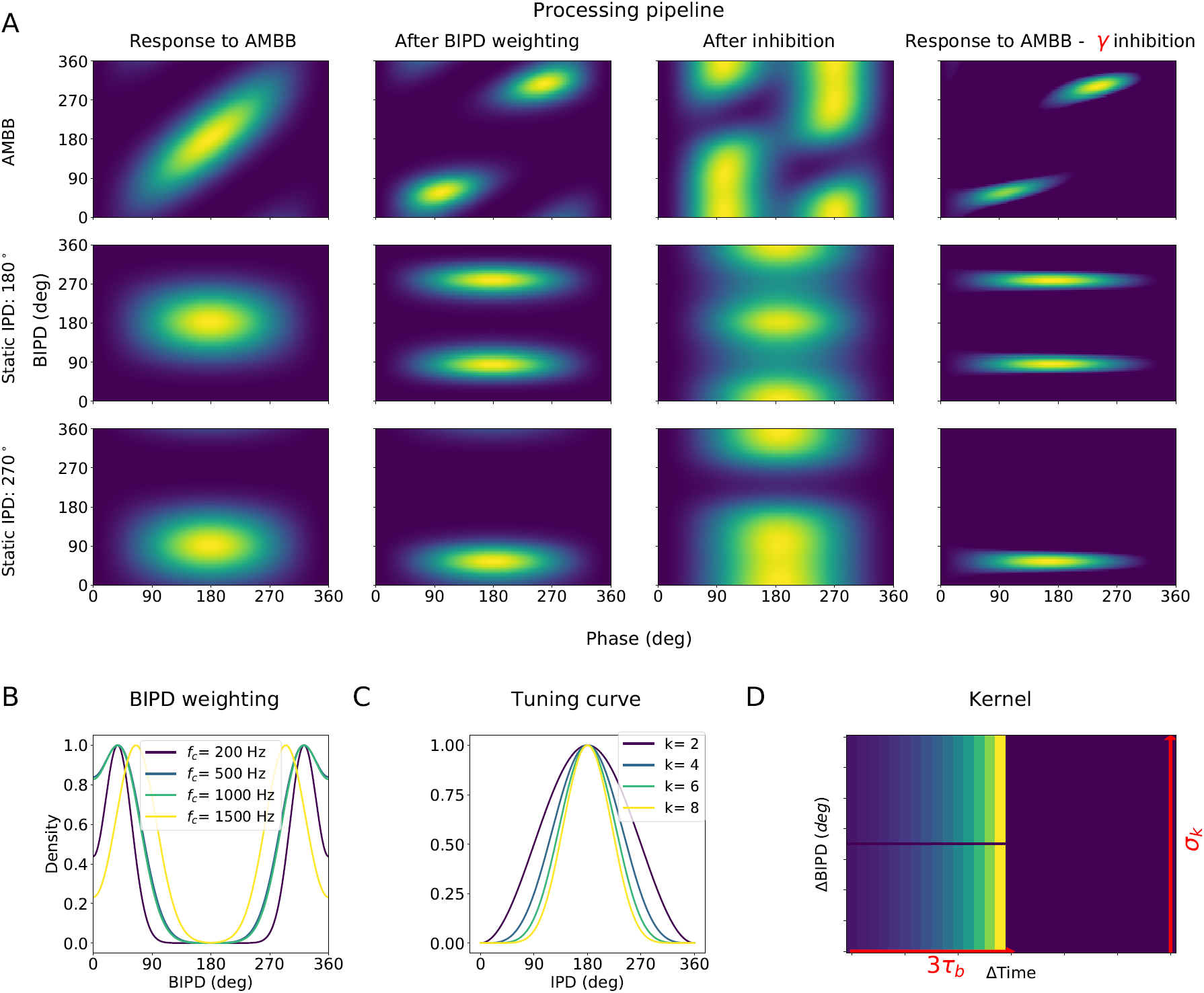
(A) Top row: the first column shows the response of the neurons to the stimulus. The response of the network after BIPD weighting is shown in the second column. The third column shows the inhibition layer, i.e. the network after inhibition. The last column shows the response of the network after subtracting the *γ* weighted inhibition layer from the BIPD weighted neural activity. Second row: Same as first row but when the stimulus is an AM tone with a static IPD=180°. Third row: same but with IPD=90°. (B) Shape of the BIPD distribution at different carrier frequencies. (C) Shape of the tuning curve centered around *BIPD* = 180° with different values of the exponent *k*. (D) Shape of the inhibition kernel. The weight of inhibition decreases exponentially with time following the time constant *τ*_*b*_. The fraction of neurons involved in the inhibition process is set by the variable *σ*_*k*_.

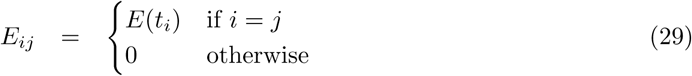

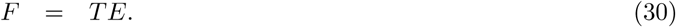

The pattern of activity generated by equation (30) can be seen in Figure 10A.

#### BIPD distribution

We used an experimentally measured BIPD distribution, with the mean *μ* and the standard deviation *σ* of the BIPDs measured from guinea pigs in McAlpine et al. (2001). The density of BIPDs in our model then followed a bimodal normal distribution based on these statistics. The density function of this distribution is proportional to:

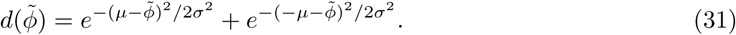

At *f*_*c*_ = 500 Hz, this distribution has values close to 0 at BIPDs around 180° and exhibits two peaks respectively around IPDs equal to 45° and 315°. BIPD distributions at different frequencies can be seen in figure 10B.

To incorporate the non-uniform BIPD distribution into the uniform matrix formulation above, we multiply the response *F*_*ij*_ with the density 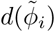 (figure 10A). The model here is continuous valued, but can be considered to represent mean firing rates or spike counts. Multiplying by the density therefore makes the response proportionate to the mean number of spikes from neurons tuned to a given BIPD. As a consequence, some of the weighted responses with 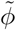 close to 180° will be close to 0 because there are next to no neurons with BIPD close to 180°. We write the response of the weighted network as *F*_*W*_, in matrix form:

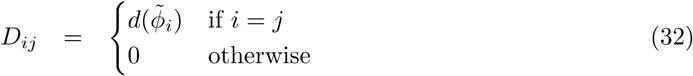

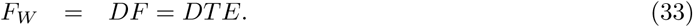

#### Lateral inhibition

The next layer of neurons in the network have a lateral inhibition mechanism. This is modelled with a synaptic connectivity assuming a narrowly tuned excitation and broad lateral inhibition (figure 11). We assume that excitatory neurons are fast compared to inhibitory neurons, which we model by applying a low-pass filter to the inhibitory neurons (approximating a number of neural processes such as membrane potential or synapse dynamics).

**Figure 11:**
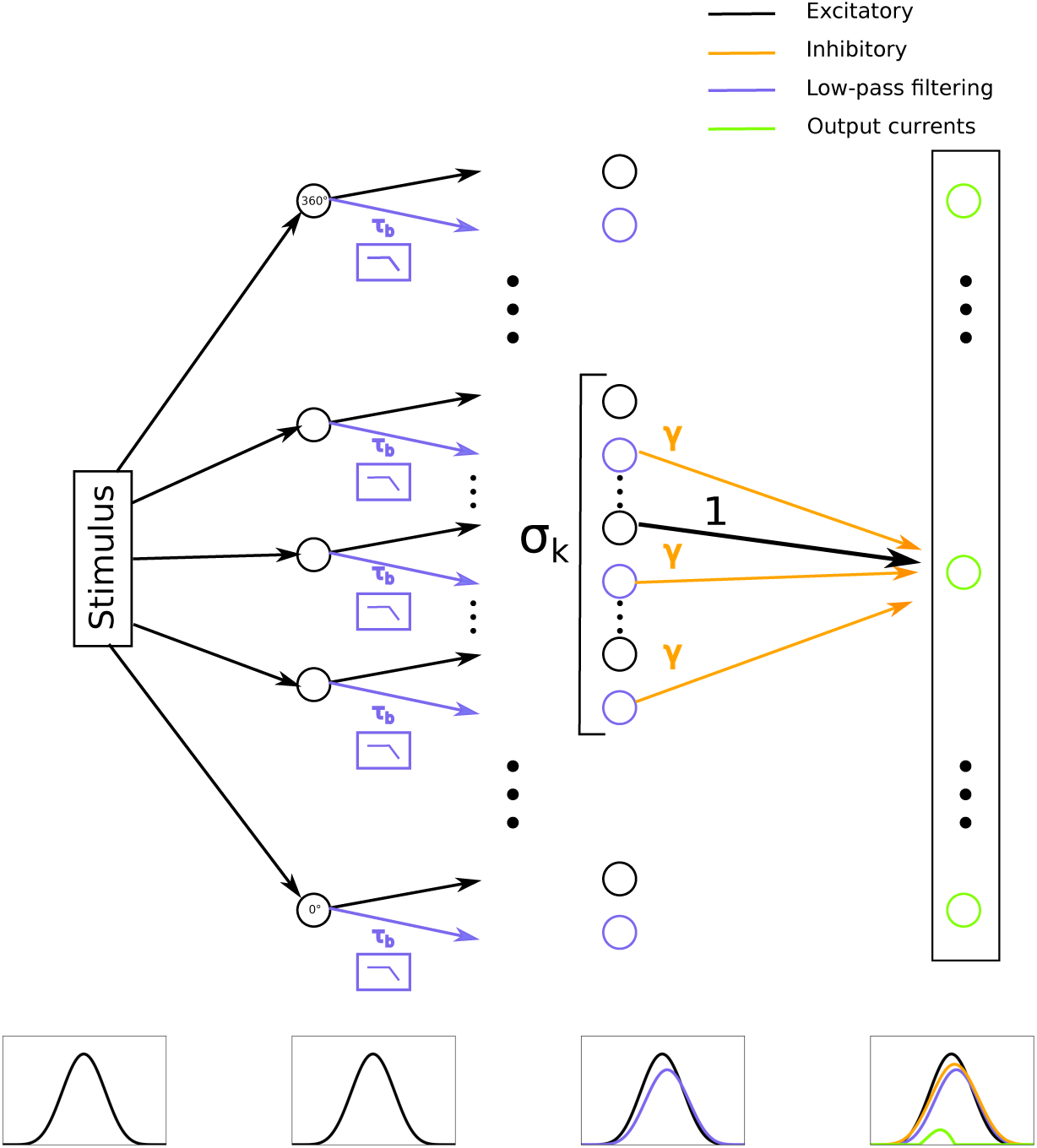
Structure of the network. Black circles represent the initial response of the neurons in the network to the AMBB stimulus. Each neuron is tuned to different BIPDs, from 0° to 360°. Blue circles represent the local inhibitory currents, and are low-pass filtered version of the black circles. The resulting inhibition currents, after low-pass filtering, are weighted by a factor *γ* and summed within the BIPD window of width *σ*_*k*_. Green circles represent the output neural activity for each BIPD. Each green circle is the response of the neuron to the stimulus (the black circles) minus the weighted sum of the inhibitory currents (orange arrows). Pattern-match decoding is performed on the pattern of currents represented by the green circles. Curves below the schematic represent the exaggerated output of the network at each stage of processing. Line colours correspond to the network elements above.

The individual inhibitory currents 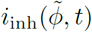 are found by low-pass filtering the response. This can be represented as a convolution of the response with an exponentially decaying kernel:

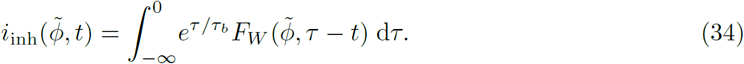

The second layer of neural responses is then formed by taking the first layer as the narrowly tuned excitatory response, and subtracting the inhibitory currents (weighted by a factor *γ*) for all BIPDs within a window of size *σ*_*k*_. The variable *σ*_*k*_ is the fraction of the set of BIPDs involved in the inhibition, so *σ*_*k*_ = 0 represents no inhibition and *σ*_*k*_ = 1 represents all neurons receiving the same inhibition proportional to the sum of the population activity. The summed inhibitory current is given by

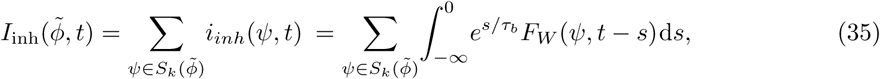

where 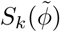 is the set of BIPDs within a distance *σ*_*k*_ of 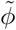 The net response of the second layer is therefore the excitatory response minus *γ* times the summed inhibitory current:

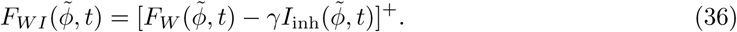

From equations 34 and 35, we can see that the summed inhibitory current *I*_inh_ can be written as the 2D convolution of the response of the network with a kernel *K*. In other words, equation (36) can be rewritten as

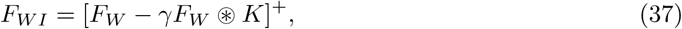

where ⊛ represents 2D convolution. The kernel *K* is the outer product of *K*_*t*_ and *K*_*ϕ*_, defined as

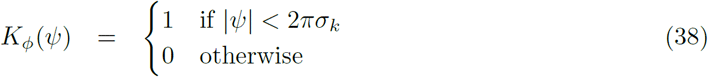

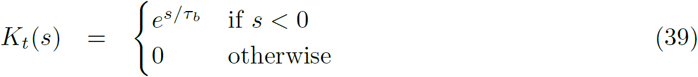

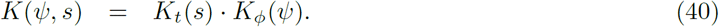

For the discretized formulation, we approximate *K* by cutting off *s <* −3*τ*_*b*_. With this approximation made, equation 37 becomes a matrix equation (although note that some care is needed to handle the circular boundaries in the convolution). This matrix formulation was essential in optimising the model to run in a reasonable time and allowing us to evaluate such a large number of parameter sets.

The result of the convolution process is shown in the fourth column of figure 10A. An important characteristic of the network’s response after lateral inhibition is the introduction of phase-shifts in the neuronal response. The distribution of these phase-shifts mostly depend on the value of *τ*_*b*_ and on the modulation frequency (figure 12A and 12B) and becomes critical during the similarity scoring process.

**Figure 12:**
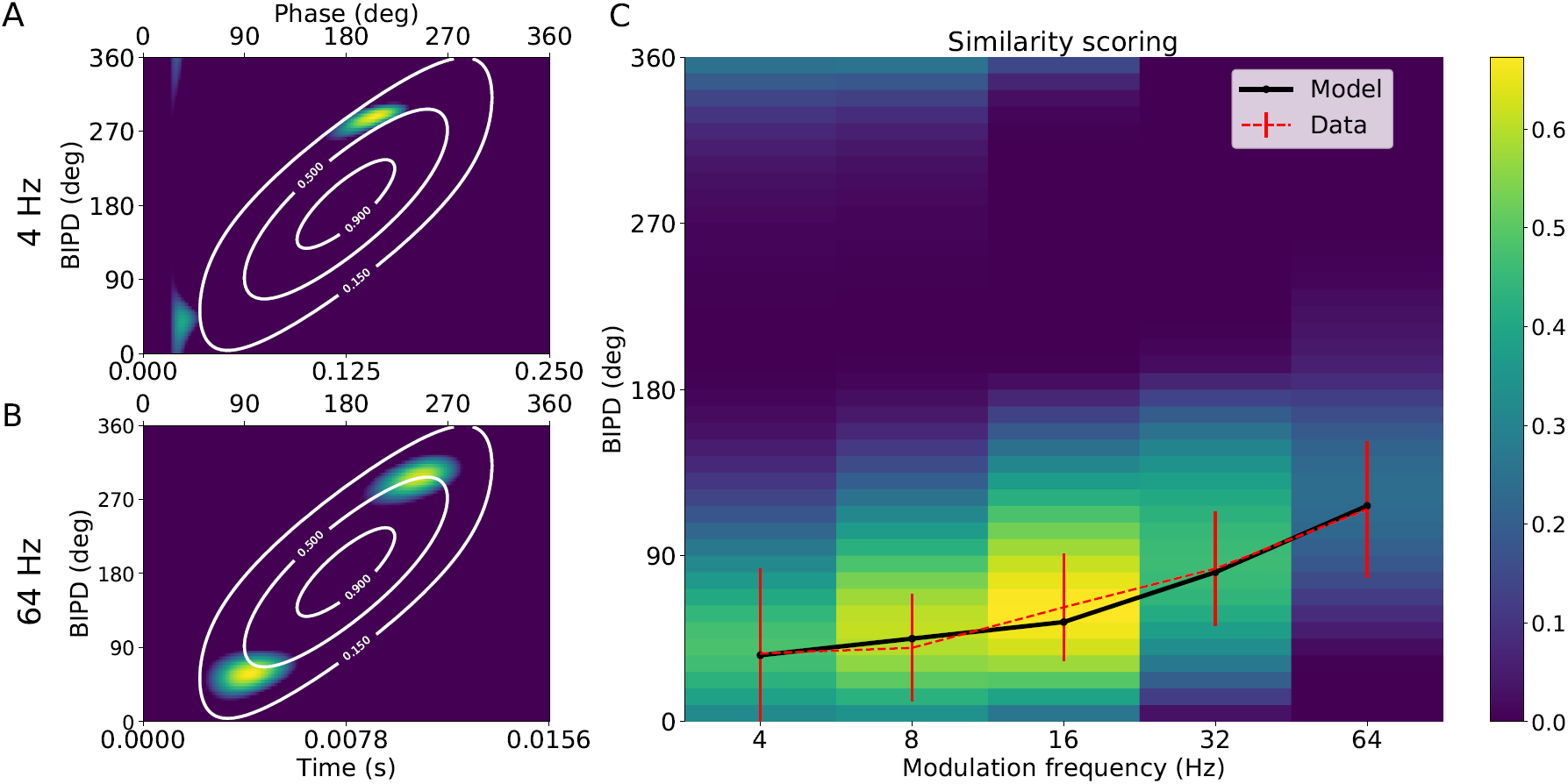
(A) Response of the network to the modelled amplitude modulated binaural beat (AMBB) stimulus with modulation frequency *f*_*m*_ = 4 Hz. Response of the network without inhibition is shown in white contours. (B) Same as A but with *f*_*m*_ = 64 Hz. (C) Similarity scoring. Similarity map showing similarity values for each modulation frequency, between the response of the full model to AMBB and the response to the amplitude modulated tones with static interaural phase differences.

#### Similarity scoring

The experiment described in Dietz et al. (2013) relies on the subjects matching static IPD pointers to the location they perceived while listening to the AMBB stimulus. To model this process, we used a similarity scoring method between the response produced by the AMBB stimulus and the response produced by different static IPD pointers. We initially considered the hemispheric decoder (Stecker et al., 2005; Grothe et al., 2010) but we were not able to use it in the context of this experiment because of its mono-dimensionality. Indeed, if the hemispheric ratio in response to the AMBB stimulus as a function of phase difference is *R*(*ϕ*) then *R* must be 2*π*-periodic by construction of the experiment. This means that all but at most two values of *R* must have multiple inverses in [0, 2*π*) and therefore it is impossible to use the hemispheric difference to match the static IPD successfully. Instead, we used a decoder inspired by the pattern match decoder described in Goodman et al. (2013). It works by computing a similarity measure between the response of the network *F*_*WI*_ to the AMBB stimulus and the response of the network to an amplitude modulated tone with a static IPD 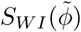. The similarity was scored over a full stimulus cycle. 40 static IPDs were selected, linearly distributed between 0° and 360°. Examples of such patterns can be found in figure 10A second and third row. We score the similarity between the response to the AMBB stimulus and the static IPD stimuli using the dot product between the flattened arrays *F*_*WI*_ and 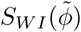 The IPD maximizing the similarity scoring was returned by the model (figure 12C). This is equivalent to the cosine similarity metric as 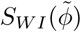 is the same for all 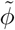

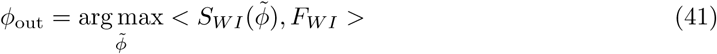

#### Starting phase

Simulations were run with 8 different starting IPDs (*ϕ*_start_) between 0° and 360° as in Dietz et al. (2013). *ϕ*_start_ corresponds to the value of the instantaneous IPD at the onset of the stimulus (when *t* = 0). In order to compare to the curves in Dietz et al. (2013), model curves were plotted using the circular mean of the extracted IPDs over the 8 different *ϕ*_start_.

## Acknowledgements

We would like to thank Mathias Dietz, David McAlpine and Andrew Brughera for fascinating discussions of this work and comments on the paper.

## Appendix

### Onset cell is a special case of excitation/inhibition model

In the single neuron onset model (equations 9-11), *R*_*e*_ could represent an excitatory pathway, *R*_*i*_ an inhibitory pathway, and *R* = [*R*_*e*_ *− βR*_*i*_]^+^ the output of excitation minus inhibition with a relative inhibitory strength *β*. However, if *β* = 1 and *τ*_*e*_ *≪ τ*_*i*_ the same equations can also be interpreted as a single onset neuron. The equations are approximately equal to a rate-based version of the octopus cell model of Spencer et al. (2018), which is itself a simplification of the models of Spencer et al. (2012) and Ferragamo and Oertel (2002). In their spiking model,

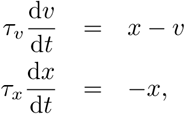

where *v* is the membrane potential, *x* is a synaptic conductance that is increased by a fixed amount for each incoming spike, and the model fires a spike when d*v/*d*t* crosses some threshold. In the limit of a large number of incoming spikes with small synaptic weights we can replace the discrete changes in *x* by a continuous current *I*(*t*) to give

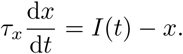

Setting *I*(*t*) = *R*_*a*_(*t*) and *τ*_*x*_ = *τ*_*e*_ we can immediately see that *R*_*e*_ = *x*. Now *v* evolves with time constant *τ*_*v*_ which is much larger than *τ*_*x*_ so on the time scale of the evolution of *v* we will have that *x ≈ I*(*t*) and so

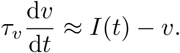

Setting *τ*_*v*_ = *τ*_*e*_ this gives us *v ≈ R*_*i*_. In the limit of a large number of output spikes, the firing rate of the onset cell will be approximately proportional to

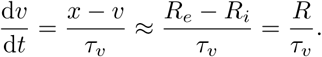

In other words, the firing rate is proportional to *R*.

### Analytic solution for *α* = 0

We set up the following set of differential equations:

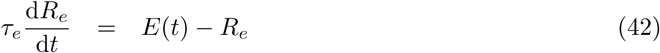

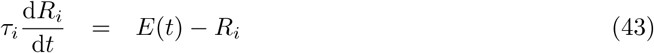

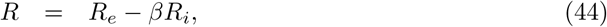

where *E*(*t*) is the envelope as above. We solve the differential equations with the following periodic boundary conditions (corresponding to the settled solution after the initial few cycles):

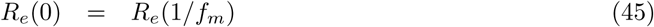

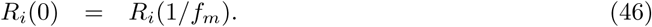

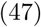

Solving these equations gives

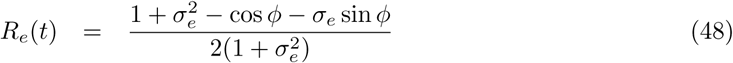

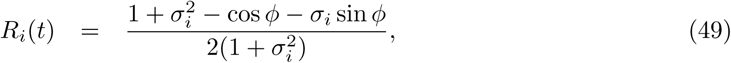

Where

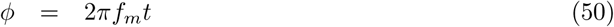

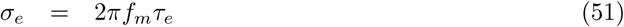

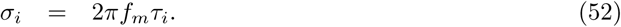

We want to find the maximum value of *t* where d*R/*d*t* = 0 and d^2^*R/*d*t*^2^ *<* 0, which we find to be the value of *θ* so that

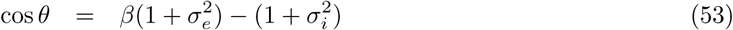

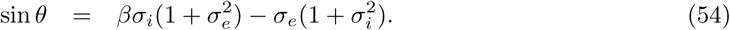

The parameter *θ* can be extracted directly as *θ* = tan^*−*1^(sin *θ/* cos *θ*) but some care has to be taken to get the correct value of *θ ∈* [0, 2*π*). This gives us a three parameter equation that can fit the data extremely well (figure 13). If we measure *f*_*m*_ in kHz then the following is a close fit to the data:

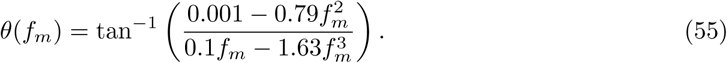

**Figure 13:**
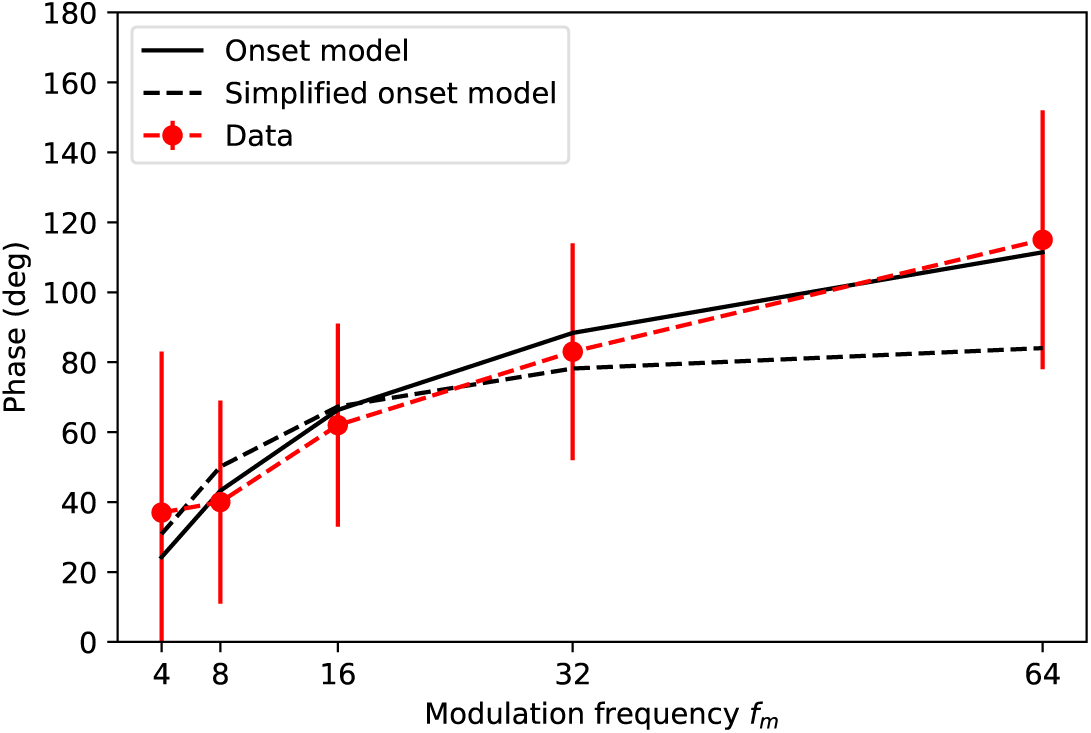
Best fit to data for analytically solved single neuron onset model.

However, we can simplify further if we assume that 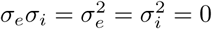 to get

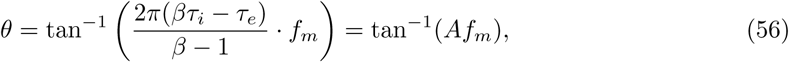

for a single constant *A*. Figure 13 shows that this approximation gives a reasonable fit with *A* = 0.15 when *f*_*m*_ is small but breaks down for *f*_*m*_ = 64 Hz.

